# Rates but not acoustic features of ultrasonic vocalizations are related to non-vocal behaviors in mouse pups

**DOI:** 10.1101/2022.08.05.503007

**Authors:** Nicole M. Pranic, Caroline Kornbrek, Chen Yang, Thomas A. Cleland, Katherine A. Tschida

## Abstract

Mouse pups produce ultrasonic vocalizations (USVs) in response to isolation from the nest (i.e., isolation USVs). Rates and acoustic features of isolation USVs change dramatically over the first two weeks of life, and there is also substantial variability in the rates and acoustic features of isolation USVs at a given postnatal age. The factors that contribute to within-age variability in isolation USVs remain largely unknown. Here, we explore the extent to which non-vocal behaviors of mouse pups relate to the within-age variability in rates and acoustic features of their USVs. We recorded non-vocal behaviors of isolated C57BL/6J mouse pups at four postnatal ages (postnatal days 5, 10, 15, and 20), measured rates of isolation USV production, and applied a combination of hand-picked acoustic feature measurements and an unsupervised machine learning-based vocal analysis method to examine USV acoustic features. When we considered different categories of non-vocal behavior, our analyses revealed that mice in all postnatal age groups produce higher rates of isolation USVs during active non-vocal behaviors than when lying still. Moreover, rates of isolation USVs are correlated with the intensity (i.e., magnitude) of non-vocal body and limb movements within a given trial. In contrast, USVs produced during different categories of non-vocal behaviors and during different intensities of non-vocal movement do not differ substantially in their acoustic features. Our findings suggest that levels of behavioral arousal contribute to within-age variability in rates, but not acoustic features, of mouse isolation USVs.

## Introduction

Following birth, mouse pups are blind, deaf, and possess limited sensorimotor capacity, relying on their mother for food and thermoregulation (Brust et al., 2015; Theiler, 1972). During their first two weeks of life, they produce vocalizations in the ultrasonic range (i.e., USVs) which are emitted in response to isolation from the dam and cold exposure (Hahn & Schanz, 2002; Okon, 1970a; Zippelius & Schleidt, 1956). These so-called isolation USVs induce a maternal retrieval response and are crucial for the survival of these altricial pups during their first two weeks of life (Ehret, 1992; Ehret & Haack, 1984; Noirot, 1972).

Both rates and acoustic features of mouse isolation USVs change over early development (Grimsley et al., 2011; Rieger & Dougherty, 2016). Rates of isolation USVs of C57 mouse pups start out low shortly after birth, peak at around postnatal day (P) 6-7, and gradually decline as pups gain the ability to thermoregulate and locomote efficiently, disappearing after 2 weeks of age (Hahn et al., 1998; Okon, 1970a; Thornton et al., 2005). With regard to developmental changes in USV acoustic features, previous studies have reported that USV duration increases over early development and then gradually declines as pups approach two weeks of age (Castellucci et al., 2018; Grimsley et al., 2011; Liu et al., 2003; Yin et al., 2016). Meanwhile, the mean pitch of isolation USVs becomes less variable over early development (Grimsley et al., 2011; Liu et al., 2003), and the interval between consecutive USVs decreases (Castellucci et al., 2018; Grimsley et al., 2011; Liu et al., 2003; Yin et al., 2016). These developmental changes in the acoustic features of isolation USVs are thought to be driven by physiological changes that take place during the first two postnatal weeks, including changes to the larynx, lung capacity, and vocal-respiratory coordination (Dutschmann et al., 2014; Elwood & Keeling, 1982; Riede et al., 2020; Wei et al., 2022).

Rates and acoustic features of isolation USVs also exhibit substantial variability within a given postnatal age, both between pups and between recordings from the same pup, even when pup and chamber temperature are monitored and maintained in a narrow range (Grimsley et al., 2011; Rieger & Dougherty, 2016). Previous studies have explored how olfactory cues (Branchi et al., 1998; D’Amato & Cabib, 1987; Ehret, 2005; Olejniczak et al., 1999; Santucci et al., 1994; Szentgyörgyi & Kapusta, 2004) and tactile cues (Branchi et al., 1998; Okon, 1970b) influence rates of isolation USV production in rodent pups. The factors that contribute to the substantial variability in rates and acoustic features of isolation USVs that occurs even in the absence of these specific environmental and social cues, however, remain unknown.

One factor that may contribute to within-age variability in the rates and acoustic features of isolation USVs is the non-vocal behavior exhibited by mouse pups (Fox, 1965). Throughout early development, motor abilities such as locomotion and grooming develop rapidly and become increasingly refined throughout the postnatal period (Brust et al., 2015; Fox, 1965). Different non-vocal behaviors might differentially influence laryngeal and vocal tract configuration and are likely associated with different rates of respiration and levels of behavioral arousal, factors which may in turn influence rates and acoustic features of USVs. To date, only a single study has examined the relationship between non-vocal behavior and variability in the rates and acoustic features of isolation USVs (Branchi et al., 2004). The authors report that P7 CD-1 Swiss mouse pups produce higher rates of isolation USVs when they engage in locomotion relative to other non-vocal behaviors (Branchi et al., 2004). However, the extent to which non-vocal pup behaviors relate to within-age variability in rates and acoustic features of isolation USVs, and how these relationships change over early development, remains poorly understood.

To address these questions, we recorded isolation USVs and non-vocal behaviors of male and female C57BL/6J mice at four postnatal ages (P5, P10, P15, and P20). At each age, we quantified rates of isolation USVs and examined USV acoustic features, using hand-picked features as well as an unsupervised machine learning-based acoustic analysis method. Categories of non-vocal behaviors were scored by trained observers, and we also quantified movement intensity from videos using an annotation and instance segmentation-based animal tracking and behavior analysis package. We then compared these descriptions of vocal and non-vocal behavior to test the hypothesis that pup non-vocal behaviors are related to within-age variability in the rates and acoustic features of isolation USVs.

## Materials and Methods

Further information and requests for resources should be directed to the corresponding author, Katherine Tschida.

### Ethics Statement

All experiments and procedures were conducted according to protocols approved by the Cornell University Institutional Animal Care and Use Committee (protocol #2020-001).

### Subjects

Male (N=21) and female (N=24) C57BL/6J mice (Jackson Laboratories, 000664) were housed with their siblings and both of their parents until weaning at postnatal day 21. Mice were kept on a 12h:12h reversed light/dark cycle and given *ad libitum* food and water for the duration of the experiment. Because we inadvertently failed to save N = 2 video recordings from P5 mice and N = 1 video recording from P10 mice, the sample sizes within each age group are as follows: P5, N = 43 mice; P10, N = 44 mice; P15, N = 45 mice; P20, N = 45 mice.

### Study design

Measurements of the vocal and non-vocal behaviors of young mice were recorded longitudinally across early development, beginning at postnatal day 5 and repeated every fifth day until postnatal day 20. Pups were identified and tracked individually by placing markings on their tails with permanent markers that were renewed every other day. For USV recordings, pups were placed in a custom acrylic chamber inside a sound-attenuating recording chamber (Med Associates) equipped with an ultrasonic microphone (Avisoft), infrared light source (Tendelux), and webcam (Logitech, with the infrared filter removed to enable video recording under infrared lighting). To elicit the production of isolation USVs, pups were recorded alone in the chamber for 5 minutes (isolation sessions). As part of a different study aimed at testing the effects of social partners on the production of isolation-induced USVs, pups were then exposed to either their mother (dam social group), a novel adult female (novel female social group), or no social partner (control group) for 5 minutes (social sessions). In the final 5 minutes, pups were again recorded alone in the chamber (re-isolation sessions). USV data from isolation and re-isolation sessions were pooled for each mouse in the current study, as we found that re-isolation USV rates did not differ between groups, nor did the acoustic features of USVs produced during isolation and re-isolation sessions differ (Fig. S1A-C). Ambient temperature inside the test chamber was maintained between 21.5-22.9° C (min/max recorded across all trials). To minimize temperature drops during transfer into the test chamber, pups were brought into the testing room within their home cage and were placed into the test chamber immediately before the start of the trial.

### USV recording, detection, and calculation of hand-picked acoustic features

USVs were recorded using an ultrasonic microphone (Avisoft, CMPA/CM16), connected to an Avisoft recording system (UltrasoundGate 116H, 250 kHz sample rate). USVs were detected using custom MATLAB codes (Tschida et al., 2019) with the following parameters implemented to detect USVs: mean frequency > 45 kHz; spectral purity > 0.3; spectral discontinuity < 1.00; minimum USV duration = 5 ms; minimum inter-syllable interval = 30 ms). For every detected USV syllable, we calculated three acoustic features: (1) duration (ms), (2) inter-syllable interval (ISI, defined as the interval from the onset of one USV to the onset of the next USV, > 400 ms intervals excluded from analyses), and (3) mean frequency (measured in kHz; dominant frequency calculated at each time point of the USV, then averaged across entire syllable).

### Analyzing USV acoustic features with Autoencoded Vocal Analysis

Isolation USVs detected with custom MATLAB codes were analyzed using Autoencoded Vocal Analysis (AVA) v0.2 (Goffinet et al., 2021), a Python package to describe and quantify animal vocalizations. Briefly, AVA uses a variational autoencoder (VAE; Kingma & Welling, 2014), an unsupervised learning method that learns from the data by training two probabilistic maps, an encoder and a decoder. Both maps are parameterized via deep convolutional neural networks. Spectrograms of individual USVs are encoded into latent representations and decoded to create reconstructions. We trained the VAE on a total of 21,622 USV syllables. We used the following pre-processing parameters to train the VAE: min_freq = 30 kHz, max_freq = 110 kHz, nperseg = 1024, noverlap = 512, spec_min_val = 3.3, spec_max_val = 7.0, mel = False, time_stretch = True, within_syll_normalize = False. Following pre-processing, n = 22 USVs that fell outside of these parameters were excluded from the final model. The multi-dimensional latent space was visualized by compressing the output of the VAE into 2-dimensional space using a uniform manifold approximation and projection (UMAP) algorithm (McInnes et al., 2020).

We quantified differences between distributions of latent syllable representations with Maximum Mean Discrepancy (MMD) (Gretton et al., 2012). To test whether USVs produced during isolation and re-isolation sessions differed in their acoustic features, we established a baseline comparison for each age group by assigning pups to one of two groups and then calculating MMD between these two distributions of isolation USVs (within session comparisons, Fig. S1B-C). We then compared these MMD values to MMD values calculated between distributions of latent syllable representations from isolation vs. re-isolation sessions within each age group (between sessions comparisons, Fig. S1C). To determine whether pups from different age groups produced USVs with different acoustic features, we first established the baseline level of acoustic variability within each age by considering USVs produced during isolation sessions vs. USVs produced during re-isolation sessions and calculating MMD between these two distributions (Fig. S1D). We then compared these within-age MMD values to MMD values quantifying the difference between distributions of latent syllable representations from pairs of different ages (P5 vs. P10, P5 vs. P15, and P10 vs. P15; Fig. S1D). Finally, to compare the acoustic features of USVs produced in each age group during different non-vocal behaviors (see next section), MMD values were calculated between distributions of latent syllable representations produced during a single category of non-vocal behavior (within behavior comparisons, pups from each age split into two groups as above) or produced during different categories of non-vocal behaviors (between behavior comparisons; Fig. S4).

### Analysis of categories of non-vocal behaviors

Trained observers scored the following behaviors from webcam videos during isolation and re-isolation sessions: (1) locomotion (movement of forelimbs and/or head while four limbs are touching the surface of the testing chamber; both locomotion (P15 and P20 mice) and attempted locomotion (P5 and P10 mice) were scored); (2) wriggling (mouse on its side or in supine position, accompanied by movement of forelimbs and/or head); (3) lying still (no visible movement); and (4) grooming (see also Supplemental Movies 1-4). Behavior scores of two trained observers were averaged (inter-rater reliability: r = .99) and used in analyses.

### Analysis of non-vocal movement intensity and comparison to USV rates

A Python software package for behavior analysis, Annolid (https://cplab.science/annolid), was used to quantify non-vocal movement intensity in each trial. Briefly, the mouse pup was detected and segmented in each video frame with an instance segmentation model trained on a custom labeled dataset, generating a mask of the pup’s shape and location in each frame. The optical flow across consecutive pairs of frames was then calculated and filtered with the pup mask to generate a vector of movement intensity, which includes head, body, and limb movements. The movement vector was smoothed with a moving average filter (span = 90 frames) and linearly interpolated such that the resulting movement vector was on the same time base as the USV rate vector for each trial. USV rates were calculated for each trial by counting the number of USVs produced in 3s-long bins. The cross-covariance between movement intensity and USV rate was then calculated for each trial using the xcov function in Matlab with the “coeff” scaling option, which normalizes each vector such that the auto-covariance at time lag 0 equals 1. Thus, each cross-covariance has a maximum possible value of 1 and a minimum possible value of -1, allowing us to pool data from different trials to compare the strength of the cross-covariance between age groups. Matched comparisons were generated for each trial by comparing movement and USVs from the first 5 minutes of recording (isolation movement vs. isolation USVs) and the final 5 minutes of recording (re-isolation movement vs. re-isolation USVs). Shuffled comparisons were generated for each trial by comparing movement and USVs from non-corresponding 5-minute recording periods (isolation movement vs. re-isolation USVs, re-isolation movement vs. isolation USVs). These matched and shuffled comparisons were averaged to generate a mean matched and mean shuffled cross-covariance for each trial. A subset of trials in which 0 USVs were recorded were excluded from these analyses (N = 2 P5 trials and N = 19 P15 trials), as cross-covariances could not be calculated for these trials. Matched and shuffled cross-covariances from multiple pups from the same age group were averaged together to obtain a mean pooled matched and a mean pooled shuffled cross-covariance for each age group. Mean pooled covariance coefficients (CCs) were calculated as the maximum value of the mean pooled matched and mean pooled shuffled cross-covariances between time lags of -5 and +5 seconds for each age group. The CC for each individual trial was then calculated by measuring the value of the single trial cross-covariance at the time lag of the mean pooled CC. To calculate the significance of the cross-covariance between movement intensity and USV rates, CCs were compared for matched comparisons and for shuffled comparisons within each age group.

### Statistical analyses

To determine whether to use parametric or non-parametric statistical tests for a given comparison, we examined the normality of the residuals for the relevant data distributions (determined by visual inspection of plots of z-scored residuals). Non-parametric tests were applied for analyses of non-normally distributed distributions. For both parametric and non-parametric comparisons, two-sided statistical comparisons were used (alpha = 0.05). All p values for pairwise comparisons represent Bonferroni-corrected values. Details of the statistical analyses used in this study are included in Table S1. No statistical methods were used to pre-determine sample size. All statistical analyses were carried out using R 4.1.0 (R Core Team, 2021) and R Studio 1.4.1717 (RStudio Team, 2022).

### Data availability

All source data generated in this study as well as custom Matlab codes for USV detection will be deposited in a digital data repository, and this section will be modified prior to publication to include the persistent DOI for the dataset. Autoencoded Vocal Analysis (Goffinet et al., 2021) is freely available online: https://github.com/jackgoffinet/autoencoded-vocal-analysis. Annolid is open-source and can be obtained from the GitHub repository https://github.com/cplab/annolid. The labeled dataset and trained instance segmentation model will also be included in the digital data repository.

## Results

### Rates and acoustic features of isolation USVs change over early postnatal development

We first measured how rates of USV production changed over early postnatal development by tracking and comparing USV rates from mouse pups recorded for 10 minutes in isolation at P5, P10, P15, and P20 (Fig. 1A-B; see Methods). We found that rates of isolation USVs increased from P5 to P10, and that P10 mice vocalized at significantly higher rates than all other ages (Fig. 1B; P5: 117.98 ± 109.68 USVs, N = 43 mice; P10: 311.77 ± 201.23 USVs, N = 44 mice; P15: 66.36 ± 86.18, N = 45 mice; P20: 8.27 ± 36.86, N = 45 mice; p < 0.01 for P10 vs. all other ages and for P5 vs. P20; one-way ANOVA followed by post-hoc tests). We conclude that the production of isolation USVs by mouse pups peaks around P10 and then gradually declines, consistent with previous studies (Castellucci et al., 2018; Grimsley et al., 2011; Liu et al., 2003; Yin et al., 2016).

**Figure 1.**
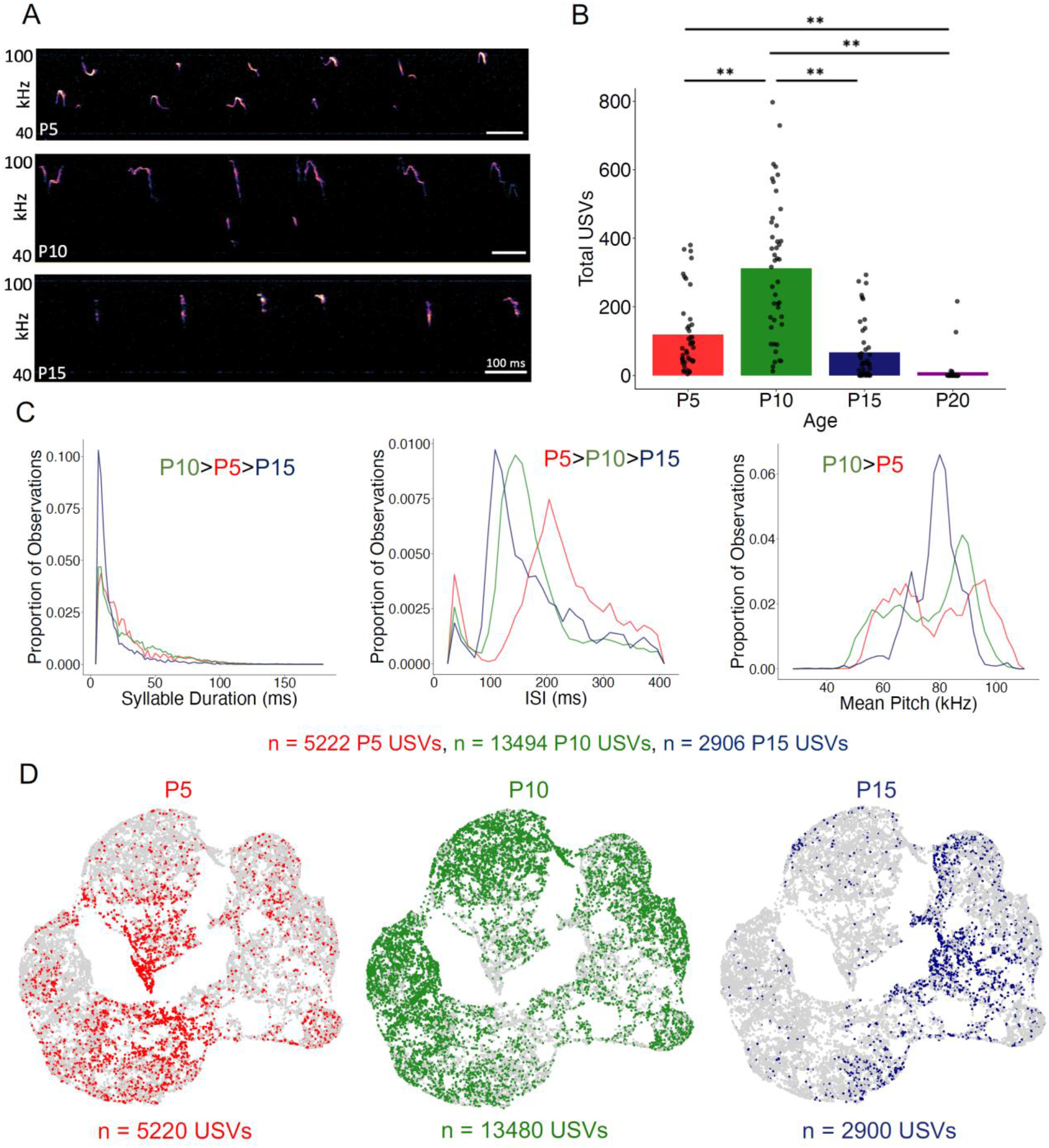
Rates and acoustic features of isolation USVs change over early postnatal development. (A) Spectrograms of representative isolation USVs produced by P5, P10, and P15 mice. (B) The number of isolation USVs is shown for P5 (N = 43), P10 (N = 44), P15 (N= 45), and P20 mice (N = 45). Double asterisks, p < 0.01. (C) (Left) Distribution of durations of isolation USVs recorded from P5 (red), P10 (green), and P15 mice (blue). (Middle) Same, for inter-syllable intervals between isolation USVs. (Right) Same, for mean pitch. Significant differences between median values of groups indicated by the text in each panel; p < 0.01 for all indicated comparisons. (D) UMAP projections of latent syllable representations of USVs produced by P5 (red), P10 (green), and P15 mice (blue). Points represent individual syllables and are closer to each other if acoustically more similar.

Next, we explored developmental changes in the acoustic features of isolation USVs produced by P5, P10, and P15 mice (P20 mice were not considered further in the study, as they produced near-zero rates of isolation USVs, Fig. 1B). Previous studies have described acoustic changes in isolation USVs over early development by quantifying hand-picked acoustic features or by examining the proportions of USVs that fall into experimenter-defined acoustic categories (Castellucci et al., 2018; Fonseca et al., 2021; Grimsley et al., 2011; Liu et al., 2003; Scattoni et al., 2008; Vogel et al., 2019; Yin et al., 2016). To assess whether the USVs recorded in our dataset exhibit similar developmental changes to those that have been previously reported, we first quantified developmental changes in USVs using a small number of hand-picked acoustic features (see Methods). Because some mice within each age group produced low rates of isolation USVs (Fig. 1B), we analyzed changes in acoustic features across development by pooling all USVs produced by mice within each age group. We found that USV durations peaked at P10 and subsequently decreased in P15 mice and that the interval between consecutive USV syllables (i.e., ISI) was highest at P5 and decreased at later ages (Fig. 1C, left and middle panels; median and IQR of duration for P5 USVs: 19.97 ± 22.53; P10 USVs: 21.50 ± 30.72; P15 USVs: 10.75 ± 11.78; p < 0.001 for all duration comparisons; median and IQR of ISIs for P5 USVs: 217.60 ± 97.79; P10 USVs: 157.18 ± 66.05; P15 USVs: 150.02 ± 105.98; p < 0.01 for all ISI comparisons; Kruskal-Wallis followed by post-hoc tests). In addition, median values of mean USV pitch were higher at P10 than at P5 (Fig. 1C, right panel; median and IQR of mean pitch for P5 USVs: 79.77 ± 27.16; P10 USVs: 79.87 ± 9.62; P15 USVs: 79.57 ± 9.61; p < 0.001; Kruskal-Wallis followed by post-hoc tests), and more notably, distributions of mean pitch changed in shape from bimodal at P5 and P10 to a narrower, more unimodal distribution for USVs produced by P15 mice (Fig. 1C, right panel). These changes in rates and acoustic features of USVs are consistent with those reported in previous studies (Castellucci et al., 2018; Grimsley et al., 2011; Liu et al., 2003; Yin et al., 2016).

A potential limitation of examining USVs using hand-picked acoustic features or syllable categories is that hand-picked features may miss important aspects of acoustic variability and may be correlated with one another, and USVs that fall within a given experimenter-defined acoustic category still exhibit substantial variability in their acoustic features (Goffinet et al., 2021; Sainburg et al., 2020). To examine developmental changes in the acoustic features of isolation USVs in a manner that does not depend on hand-picked acoustic features or clustering of USVs into categories, we examined and compared vocal repertoires of mice using Autoencoded Vocal Analysis v0.2 (AVA; Goffinet et al., 2021), a recently described unsupervised modeling approach that uses VAEs (Kingma & Welling, 2014). In this method, VAEs use spectrograms of individual vocalizations as inputs, apply an encoder to represent and compress these spectrograms into a small number of latent features, and then use a decoder to reconstruct the input spectrograms (see Fig. S2A for reconstructed spectrograms of representative syllables). We provided spectrograms of isolation USV syllables produced by P5-P15 mice as input to train the model and found that the VAE converged on a latent representation of four dimensions. We then compressed the resulting latent features with a dimensionality reduction method, uniform manifold approximation and projection (UMAP; McInnes et al., 2020) and visualized latent syllable representations in 2D space with Bokeh plots. These plots revealed that USVs of P5 and P15 mice were distributed in a largely non-overlapping manner, while P10 USVs were distributed relatively evenly across the acoustic space (Fig. 1D; see also Fig. S2 for UMAP representations of USV syllables color-coded by hand-picked acoustic features).

To quantify differences between these distributions, we calculated the Maximum Mean Discrepancy (MMD) between distributions of latent syllable representations to generate three comparisons: P5 vs. P10 USVs, P5 vs. P15 USVs, and P10 vs. P15 USVs. In addition, the baseline level of acoustic variability within each age was estimated by calculating MMDs between USVs recorded during the first 5 minutes to those recorded during the final 5 minutes of isolation within each age group (Fig. S1D). These comparisons revealed clear differences in vocal repertoires between P5, P10, and P15 mice, in agreement with our analyses of hand-picked features, and provide us with a rich and comprehensive description of acoustic changes across early development that we used subsequently to interrogate the relationship between USV acoustic features and non-vocal behaviors.

### Non-vocal behaviors of mouse pups change during early development

To begin to understand how USV rates and acoustic features relate to non-vocal behaviors, we examined how non-vocal behaviors of isolated mice changed from P5 to P15. We recorded four different behaviors produced by mouse pups: grooming, wriggling (movement of forelimbs and/or head while pup is on its side or in supine position), locomotion/attempted locomotion, and lying still (see Methods and Supplemental Movies 1-4). Unsurprisingly, we found that the rates at which mouse pups produce different non-vocal behaviors changes over early development (Fig. 2; two-way ANOVA followed by post-hoc tests). Specifically, we found that P5 mice spend more time lying still and less time locomoting than P10 and P15 mice (proportion time spent at P5 lying still: 0.74 ± 0.10; P10 lying still: 0.57 ± 0.09; P15 lying still: 0.51 ± 0.20; P5 locomotion: 0.16 ± 0.12; P10 locomotion: 0.43 ± 0.09; P15 locomotion: 0.37 ± 0.16; p < 0.01 for all comparisons). Wriggling occurred most frequently in P5 mice, declined in frequency in P10 mice, and was no longer observed in P15 animals (proportion time spent wriggling at P5: 0.10 ± 0.08; P10 wriggling: 0.002 ± 0.01; p < 0.01 for P5 vs. P10 and P15 wriggling). Grooming was observed only in P15 mice (proportion time spent grooming at P15: 0.12 ± 0.08; p < 0.01 for P15 vs. P5 and P10 grooming). In summary, time spent locomoting increases and time spent lying still declines over early development, wriggling was observed mainly in P5 mice, and grooming was observed only in P15 mice.

**Figure 2.**
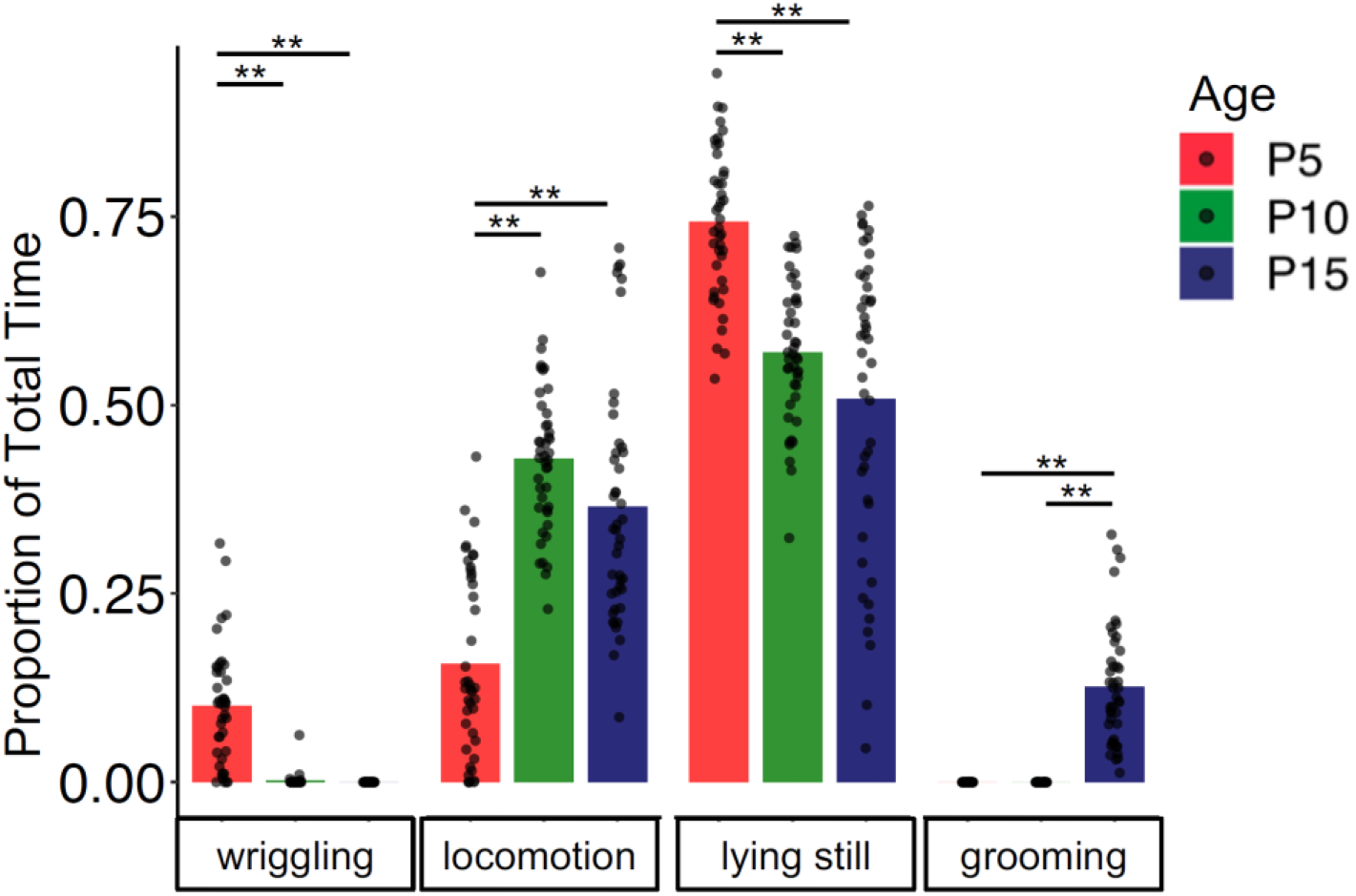
Non-vocal behaviors change over early postnatal development. The proportion of time spent engaged in different non-vocal behaviors is shown for P5 (red), P10 (green), and P15 mice (blue). Double asterisks, p < 0.01.

### Relationship of USV rates to categories of non-vocal behavior

We next investigated the relationship between USV rates and categories of non-vocal behaviors of isolated mice. In other words, are mouse pups more likely to produce USVs during certain non-vocal behaviors compared to others, and do these relationships change with age? First, we assigned a behavior code to each USV syllable that corresponds to the non-vocal behavior the mouse was performing when producing that syllable (locomotion, lying still, wriggling, or grooming; see Fig. S3 for ethograms of USV rates aligned with categories of non-vocal behavior for representative trials). Because there is substantial variability both within ages and across ages in the proportion of time that mice spend engaged in different non-vocal behaviors (Fig. 2), we then calculated the numbers of USVs produced by each mouse during different non-vocal behaviors as a function of the total time spent that mouse engaged in each non-vocal behavior (USVs per second of behavior; Fig. 3A). This analysis revealed that P5 mice produced the highest normalized rates of USVs during wriggling, followed by locomotion (mean USVs/second at P5 for locomotion: 0.61 ± 1.01; lying still: 0.21 ± 0.22; wriggling: 0.87 ± 1.20; p < 0.05 for the comparison between locomotion vs. lying still and lying still vs. wriggling; Friedman test followed by post-hoc tests). Both P10 mice and P15 mice produced the highest normalized rates of USVs during locomotion (mean USVs/second at P10 for locomotion: 2.09 ± 1.49; lying still: 0.39 ± 0.37; wriggling: 0.03 ± 0.17; p < 0.001 for all comparisons at P10; mean USVs/second at P15 for locomotion: 0.60 ± 0.56; lying still: 0.10 ± 0.15; grooming: 0.23 ± 0.28; p < 0.001 for comparisons between locomotion vs. grooming and locomotion vs. lying still at P15; Friedman test followed by post-hoc tests). P15 mice also produced higher normalized rates of USVs during grooming than during lying still (mean USVs/second for grooming: 0.23 ± 0.28; lying still: 0.10 ± 0.15; p <0.05; Friedman test followed by post-hoc tests). We conclude that, when considering the amount of time mice spent engaged in different non-vocal behaviors, mice in all age groups produce the highest rates of USVs during active behaviors (locomotion, wriggling, and grooming) and are less likely to vocalize while lying still (Fig. 3B).

**Figure 3.**
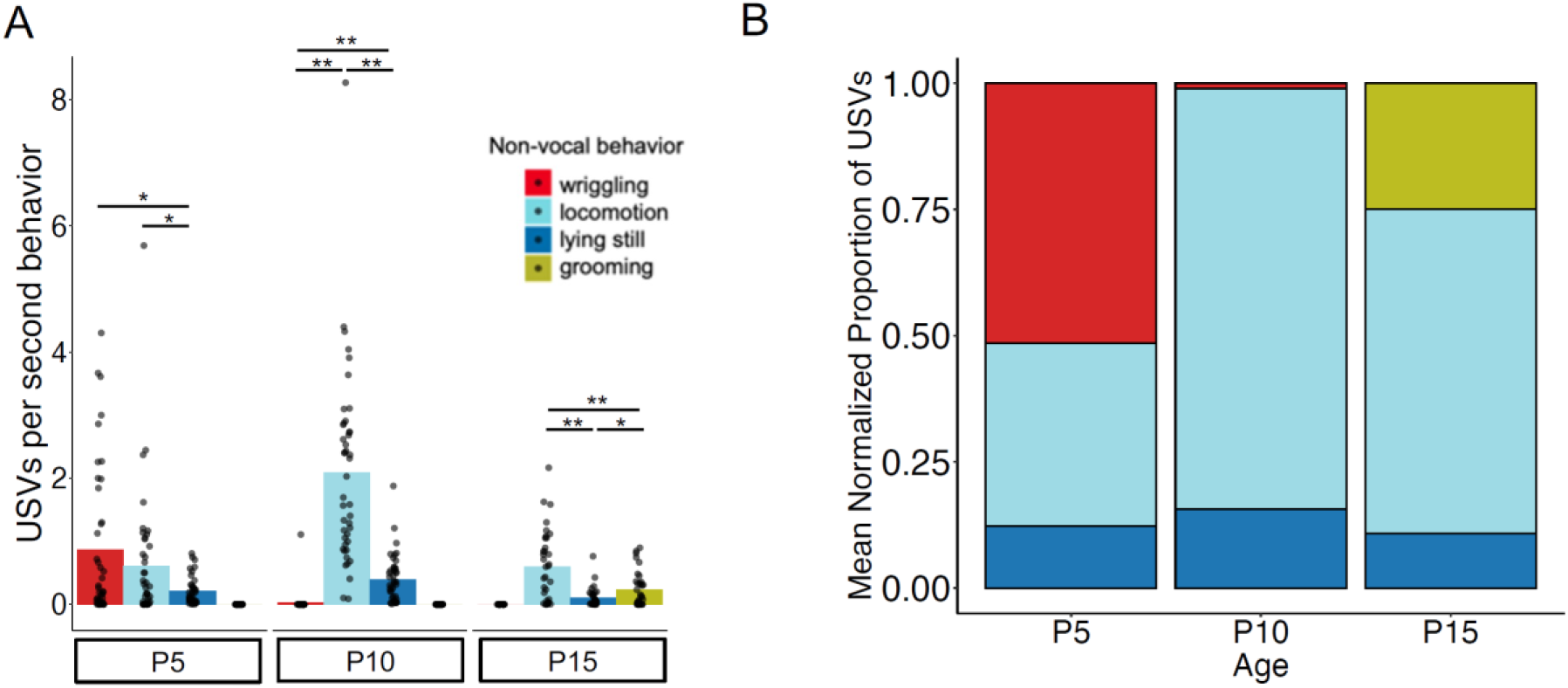
Relationship of USV rates to non-vocal behaviors. (A) The mean number of isolation USVs produced per second engaged in four non-vocal behaviors (wriggling, locomotion, lying still, and grooming; see Methods) is shown for P5, P10, and P15 mice. (B) Stacked bar plot showing the mean proportion of total USVs produced during each non-vocal behavior normalized by the amount of time each mouse spent performing that behavior. Single asterisks, p < 0.05; double asterisks, p < 0.01.

### Relationship of USV acoustic features to categories of non-vocal behavior

We next asked whether mouse pups produce isolation USVs with different acoustic features as they engage in different categories of non-vocal behaviors. We first examined whether USVs produced during different non-vocal behaviors differ in terms of hand-picked acoustic features (Figs. 4A-C). These analyses revealed significant but small differences in the duration of USVs produced during different non-vocal behaviors within each age (Fig. 4B; comparisons with significant differences indicated by colored text within each panel, p < 0.05, Kruskal Wallis with post-hoc tests). Similarly, we found significant but small differences in the mean pitch of USVs produced during different non-vocal behaviors within each age group (Fig. 4C, p < 0.05, Kruskal Wallis with post-hoc tests). Notably, there was substantial variability in the duration and mean pitch of USVs produced during a given non-vocal behavior within each age group, and the distributions of hand-picked acoustic features of USVs produced during different non-vocal behaviors were highly overlapping.

**Figure 4.**
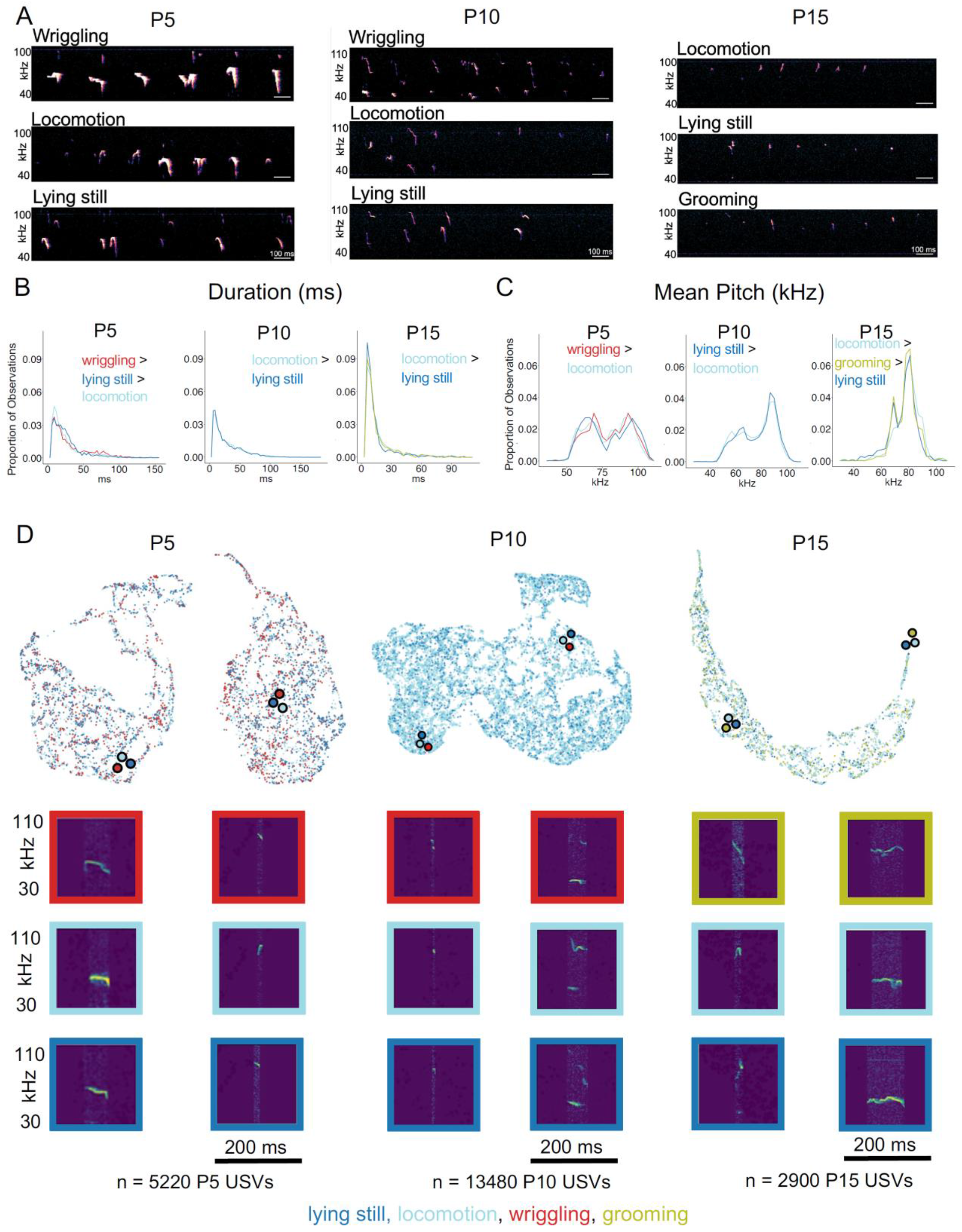
Relationship of USV acoustic features to non-vocal behaviors. (A) Spectrograms of representative isolation USVs produced by P5 (left), P10 (middle), and P15 mice (right) during different non-vocal behaviors. (B) The distributions of durations of isolation USVs produced during different non-vocal behaviors is shown for P5 (left), P10 (middle), and P15 mice (right). P10 mice only produced < 50 total USVs during wriggling, so these are excluded from analysis. (C) The distributions of mean pitch of isolation USVs produced during different non-vocal behaviors is shown for P5 (left), P10 (middle), and P15 mice (right). Significant differences between groups indicated in each panel; p values less than or equal to 0.05 for indicated comparisons. (D) UMAP projections of latent descriptions of USV syllables produced by P5 (left), P10 (middle), and P15 mice (right). Representations of individual syllables are outlined in color according to the non-vocal behavior that occurred while that syllable was produced: lying still (dark blue), locomotion (light blue), wriggling (red), and grooming (chartreuse). Example spectrograms are depicted below, and color-coded dots with black outlines indicate landmarks on the Bokeh plot for the representative spectrograms. Number of USVs at P5 produced during wriggling: 1,409, lying still: 2,017, locomotion: 1,489; P10 USVs produced during wriggling: 41, locomotion: 10,410, lying still: 2,912; P15 USVs produced during locomotion: 1,915, lying still: 478, grooming: 510.

Next, we examined and compared the acoustic features of USVs produced during different non-vocal behaviors within each age group using AVA. We assigned a non-vocal behavior code to each USV syllable and color-coded latent syllable representations according to the non-vocal behavior performed during the production of each USV (Fig. 4D; lying still USVs, dark blue; locomotion USVs, light blue; wriggling USVs, red; grooming USVs, chartreuse). This analysis revealed substantial overlap in the latent syllable representations of USVs produced during different non-vocal behaviors within each age group. To quantify differences between these distributions, we calculated the MMD between distributions of latent syllable representations to generate within-age comparisons of USVs produced during different non-vocal behaviors (Fig. S4). MMD comparisons revealed that acoustic differences between USVs produced during different non-vocal behaviors tended to be either similar in magnitude or smaller than acoustic differences between USVs produced during the same non-vocal behaviors. These analyses, together with analyses of hand-picked acoustic features, support the conclusion that mouse pups do not produce acoustically distinct types of USVs during different categories of non-vocal behaviors.

### Relationship of USV rates to non-vocal movement intensity

Although we found that rates of isolation USVs are related to categories of non-vocal behavior, these relationships were relatively weak (high variability in mean USVs per second of behavior for different pups, Fig. 3A). Even within a given category of behavior, non-vocal movements are dynamic and vary in intensity from moment-to-moment, and we wondered whether dynamic variations in movement intensity might be related to dynamic variations in USV rate. To test this idea, movement intensity in each trial was quantified using Annolid software as the total movement in the instance mask of the pup across consecutive pairs of frames (includes locomotion as well as head and limb movements, see Methods; Fig. 5A). We then calculated the normalized cross-covariance between movement intensity and USV rate for each trial (see Methods). Because each pup was recorded for 10 minutes, we separately considered the first 5 minutes and the second 5 minutes of recording to generate both matched and shuffled comparisons for each trial (Fig. 5B-C; matched: first 5 min. USVs vs. first 5 min. movement, last 5 min. USVs vs. last 5 min. movement; shuffled: first 5 min. USVs vs. last 5 min. movement, last 5 min. USVs vs. first 5 min. movement). Mean matched and shuffled cross-covariances were pooled from pups of the same age to generate mean pooled comparisons for each age group (Fig. 5C). To compare the strength of this covariance across and within different age groups, we then calculated mean pooled covariance coefficients (CC, see Methods) for matched and shuffled comparisons within each age. CCs were then calculated and compared for each trial’s matched and shuffled comparisons (see Methods; Fig. 5B). CCs for matched comparisons were significantly higher than CCs for shuffled comparisons at all ages (Fig 5B; mean matched CC’s for P5: 0.29 ± 0.21; P10: 0.42 ± 0.17; P15: 0.15 ± 0.15; mean shuffled CC’s for P5: 0.01 ± 0.07; P10: 0.00 ± 0.09; P15: 0.03 ± 0.13; p < 0.005 for all ages, two-way ANOVA with repeated measures on one factor followed by post-hoc tests). We conclude that rates of isolation USVs are well-related on average to the intensity of non-vocal movements within individual trials, and this relationship is particularly pronounced at earlier postnatal ages.

**Figure 5.**
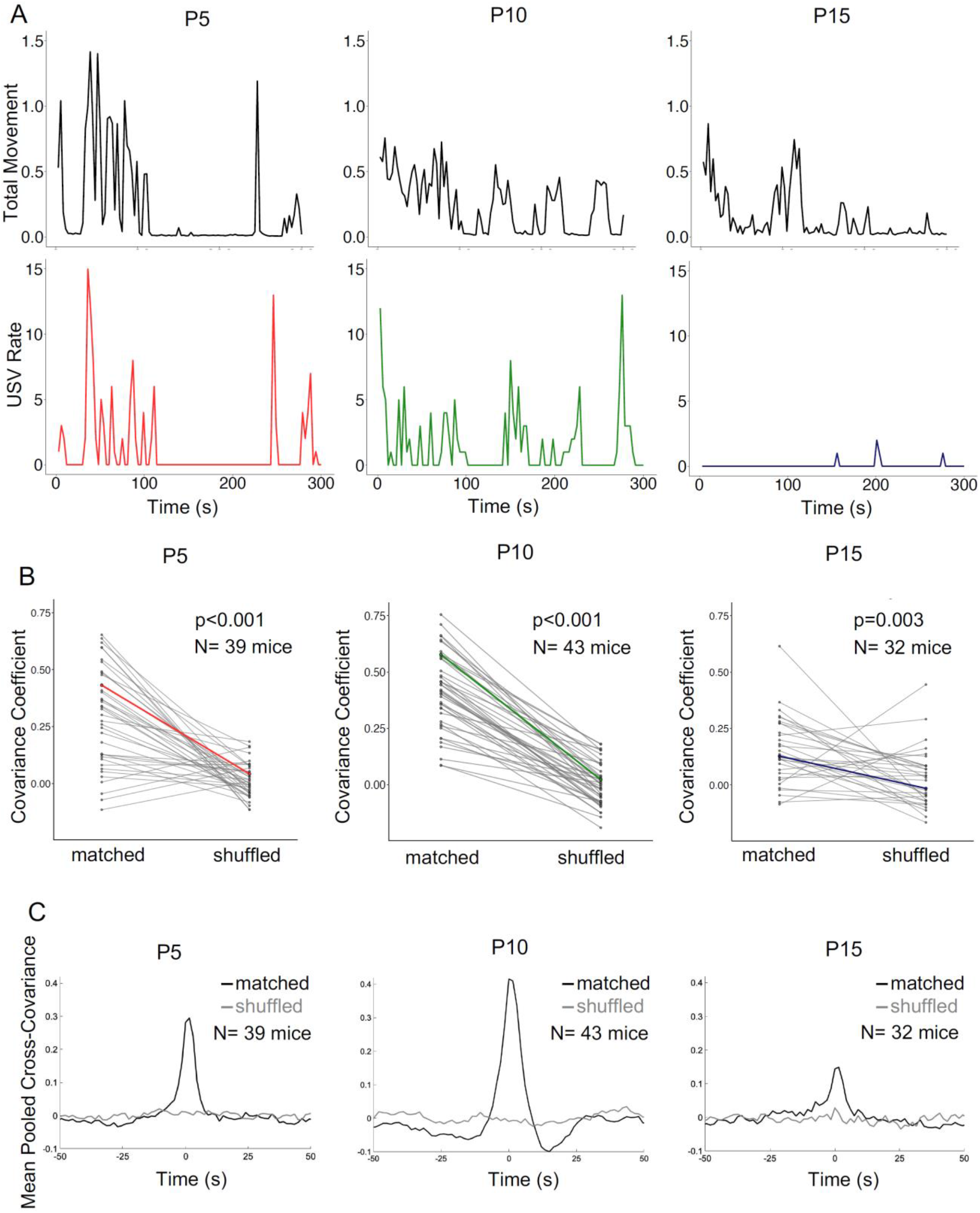
Relationship of USV rates to intensity of non-vocal movements. (A) Plots of non-vocal movement intensity (top row) and USV rate (bottom row) are shown for three representative trials. P5, left; P10, middle, P15, right. (B) Covariance coefficients for matched and shuffled comparisons are plotted for each mouse in each postnatal age group. Colored lines correspond to the examples shown in (A). (C) Mean pooled matched (black) and shuffled (gray) cross-covariances are shown for the comparison of movement intensity and USV rate in the three postnatal age groups.

### Relationship of USV acoustic features to non-vocal movement intensity

Finally, we considered whether mice produce USVs with different acoustic features as they engage in movements of different intensity. We note that USVs produced while lying still were highly overlapping in their acoustic features with USVs produced during active behaviors within each age group (Fig. 4), suggesting that USV acoustic features are unlikely to be well related to the intensity of non-vocal movements. Nonetheless, we formally considered this idea by color-coding latent syllable representations in AVA according to intensity of non-vocal movements that were ongoing at the time a given USV was produced (Fig. S5). As expected, this analysis revealed substantial overlap in the latent syllable representations of USVs produced during different intensities of non-vocal movements within each age group, indicating that non-vocal movement intensity cannot account for within-age variability in the acoustic features of isolation USVs.

## Discussion

In this study, we compared the vocal and non-vocal behaviors of mouse pups over early postnatal development to test the hypothesis that within-age variability in the rates and acoustic features of isolation USVs is related to the production of non-vocal behaviors. Consistent with previous studies (Castellucci et al., 2018; Grimsley et al., 2011; Liu et al., 2003; Yin et al., 2016), we found that rates of isolation USVs peak in P10 pups and then subsequently decline, paralleling the gradual increase in pups’ ability to locomote and thermoregulate independently (Theiler, 1972). Similarly, measurements of hand-picked features and machine-learning-based analyses of USV acoustic features revealed dramatic changes over early postnatal development, consistent with previous studies (Castellucci et al., 2018; Grimsley et al., 2011; Liu et al., 2003; Yin et al., 2016). By comparing USV rates to the production of non-vocal behaviors within each trial, we determined that pups in all age groups (P5, P10, and P15) produce higher rates of isolation USVs during active behaviors compared to periods of inactivity. Moreover, rates of USV production were well-correlated with movement intensity in individual trials, particularly in P5 and P10 pups. In contrast, the acoustic features of isolation USVs were unrelated to the production of different categories and intensities of non-vocal movements, and this was true across all age groups. To our knowledge, this is the first study to investigate the possibility that rates and acoustic features of mouse pup isolation USVs may be related to non-vocal behaviors.

Why are pups more likely to produce isolation USVs during active behaviors than during periods of inactivity and particularly during high intensity movements? One factor contributing to these relationships could be changes in respiration associated with movement. As they transition from rest to vigorous movement, animals increase their respiratory rate (Bramble & Carrier, 1983; DiMarco et al., 1983; Gravel et al., 2007; Hérent et al., 2020; Mateika & Duffin, 1995) and switch from passive to active expiration (Abdala et al., 2009; Abraham et al., 2002), a process which recruits abdominal muscles to increase intrathoracic pressure. Although no study to our knowledge has measured respiratory rates or patterns during the production of different non-vocal behaviors in mouse pups, it is possible that movement-associated changes in respiration promote USV production. On the one hand, brainstem respiratory circuits (Del Negro et al., 2018; Dutschmann & Dick, 2012; Pagliardini et al., 2011; Yackle et al., 2017; Zuperku et al., 2017; Ikeda et al., 2017) might influence the activity of brainstem neurons known to be important for USV production in adults (Hartmann & Brecht, 2020; JuÈrgens, 2002; Tschida et al., 2019) and pups (Hernandez-Miranda et al., 2017; Wei et al., 2022). A related possibility is that neuronal circuits important for movement generation, some of which in turn contribute to movement-related changes in respiration (Eldridge et al., 1981; Gariépy et al., 2012), might act on brainstem vocal-respiratory circuits to promote pup USV production.

Movement could also promote USV production as a consequence of increased intrathoracic pressure during periods of active expiration. Some studies have suggested that pup USVs are produced as a passive, acoustic byproduct of laryngeal braking, a respiratory mechanism in which contraction of abdominal muscles and constriction of the larynx improves gas exchange in the lungs during periods of high metabolic demand (Blumberg & Alberts, 1990; Blumberg & Sokoloff, 2001; Gilfoil et al., 1959; Youmans et al., 1963; see also Shair & Jasper, 2003). If laryngeal braking accounts for pup USV production in at least some instances, any factor that increases intrathoracic pressure, including the transition to active expiration during high intensity movements, might promote USV production. Finally, we note that if for any reason a pup is in a behavioral state that is favorable to USV production, movement-related increases in respiratory rates per se could lead to increased rates of USV production. Rodents produce USVs as they exhale (Roberts, 1972; Sirotin et al., 2014; Wei et al., 2022), and movement-related increases in the number of respiratory cycles per second would provide more opportunities for USV production to occur than when pups are at rest.

A second, non-mutually exclusive possibility is that movement-associated increases in arousal contribute to the observed relationships between non-vocal movements and USV rates. Previous studies have found that marmosets are more likely to produce contact calls during periods of heightened arousal, as measured by fluctuations in heart rate (Borjon et al., 2016; Gustison et al., 2019). Similarly, humans exhibit increases in heart rate and blood pressure prior to speech onset (Linden, 1987; Lynch et al., 1980). We did not include any physiological measures of arousal in the current study, but such measures could be included in future work to better understand how these variables relate to pup USV production and to what extent they can account for the relationship of non-vocal movements to pup USV rates within trials, across trials, and across early postnatal development.

While rates of isolation USVs were on average well-related to movement intensity in P5 and P10 pups, the strength of this relationship decreased in P15 mice (Fig. 5). This finding is perhaps not surprising, given that rates of isolation USVs decline as pups gain the ability to thermoregulate (Hahn et al., 1997; Okon, 1970a; Thornton et al., 2005), with isolation USV rates reaching near-zero around or shortly after two weeks of age (Fig. 1A; Castellucci et al., 2018; Grimsley et al., 2011; Liu et al., 2003; Yin et al., 2016). Interestingly, a strong relationship between USV rates and non-vocal movements re-emerges in adult rodents. Specifically, rates of 50 kHz USVs produced by adult rats are tightly coupled at the sub-second timescale to locomotion speed, both during and outside of social interactions (Laplagne & Costa, 2016). Conversely, adult rats produce 22 kHz USVs almost exclusively while immobile (Laplagne & Costa, 2016; Wöhr et al., 2005; Wöhr & Schwarting, 2008). Although adult mice produce only low rates of USVs in the absence of social partners or cues, adult males and females increase production of 70 kHz USVs during chases (Neunuebel et al., 2015; Sangiamo et al., 2020). Similar to what has been described in adult rodents (Laplagne & Costa, 2016), however, we found that the relationship between non-vocal movements and pup USV production is not obligatory. That is to say that pups can produce USVs while lying still and conversely, pups can produce non-vocal movements without vocalizing (Figs. 3, 4, 5A, S3). In addition, we note that there is trial-to-trial variability in the strength of the relationship between isolation USV rates and movement intensity (Fig. 5B). Taken together, our findings indicate that rates of isolation USVs are related to both the category and intensity of non-vocal movements, but additional factors must also contribute to within-age variability in pup USV production. These factors are likely to be state-like variables rather than stable traits, given that rates of USVs produced by an individual pup are variable across multiple same-day recordings (Barnes et al., 2017) and recordings performed on consecutive days (Rieger & Dougherty, 2016).

Although we uncovered clear relationships between non-vocal behaviors and USV rates, no such relationships were revealed in our comparisons of non-vocal behaviors and USV acoustic features. Previous studies in adult rodents have suggested that there are differences in the acoustic features of USVs produced during different non-vocal behaviors. For example, adult male mice produce USVs that are longer, lower frequency, and more harmonic during mounting as compared to USVs produced during social investigation (Hanson & Hurley, 2012; Matsumoto & Okanoya, 2016). In addition, adult male mice produce USVs that differ in acoustic features while interacting with social cues (e.g., urine) as compared to an anesthetized or live social partner (Chabout et al., 2015) and when interacting with a social partner as compared to after the social partner departs (Hanson & Hurley, 2012; Yang et al., 2013). Although not all of these studies explicitly considered or quantified non-vocal behaviors, undoubtedly, mice engage in different types and intensities of non-vocal movements in these different social contexts. In addition, a previous study concluded that acoustic features of prairie vole USVs covary with heart rate (Stewart et al., 2015), although we note that the conclusions of this study were based on only 65 USVs produced by 3 voles.

Nonetheless, in the current study, we considered many thousands of USVs produced by 45 mouse pups and found that USVs produced during different categories of non-vocal behavior did not differ substantially in their acoustic features, nor did USV acoustic features vary according to the intensity of non-vocal movements produced concurrently with vocalization. Although we found subtle differences in distributions of the duration and mean pitch of USVs produced during different non-vocal behaviors, distributions of these hand-picked acoustic features demonstrated considerable variability and were largely overlapping for USVs produced during different non-vocal behaviors (Fig. 4A-C). While most previous studies have considered USV acoustic variability by placing USVs into categories defined either by the experimenter or with the help of machine learning (Burkett et al., 2015; Coffey et al., 2019; Fonseca et al., 2021; Holy & Guo, 2005; Van Segbroeck et al., 2017; Vogel et al., 2019, 2019; Warren et al., 2021). Our findings with AVA further support the conclusion that mouse pups do not produce acoustically distinct types of USVs during different categories of non-vocal behaviors (Fig. 4D) or during different intensities of non-vocal movements (Fig. S5). Ultimately, the acoustic features of a given USV depend on respiratory rate, subglottal pressure and air flow, laryngeal muscle activation, and vocal tract configuration (Håkansson et al., 2022; Riede, 2011, 2013, 2018). Thus, our findings suggest patterns of activity in the premotor and motor neurons that control the vocal actuators are not highly constrained by pup non-vocal behavior. The factors that contribute to variability in the acoustic features of pup and adult USVs, in terms of both moment-to-moment variability within a given behavioral context as well as differences across contexts, are an important topic that warrants further study.

## Acknowledgements

Thanks to Michael Sheehan and Nora Prior for providing feedback on draft versions of this manuscript, and thanks to Jack Goffinet for help implementing AVA. Thanks also to Frank Drake and other CARE staff for their excellent mouse husbandry.

## Author contributions

N.M.P. and K.A.T. designed the experiments. N.M.P. conducted the experiments. N.M.P., C.K., C.Y., T.A.C., and K.A.T. analyzed the data. N.M.P. and K.A.T. wrote the manuscript, and all authors approved the final version.

## Notes

### Competing Interest Statement

The authors have declared no competing interest.

