## Supplemental Information for "Rates but not acoustic features of ultrasonic vocalizations are related to non-vocal behaviors in mouse pups"

### Supporting Information

**Figures S1-6**

**Movies S1-4**

**Table S1**

**Figure S1**

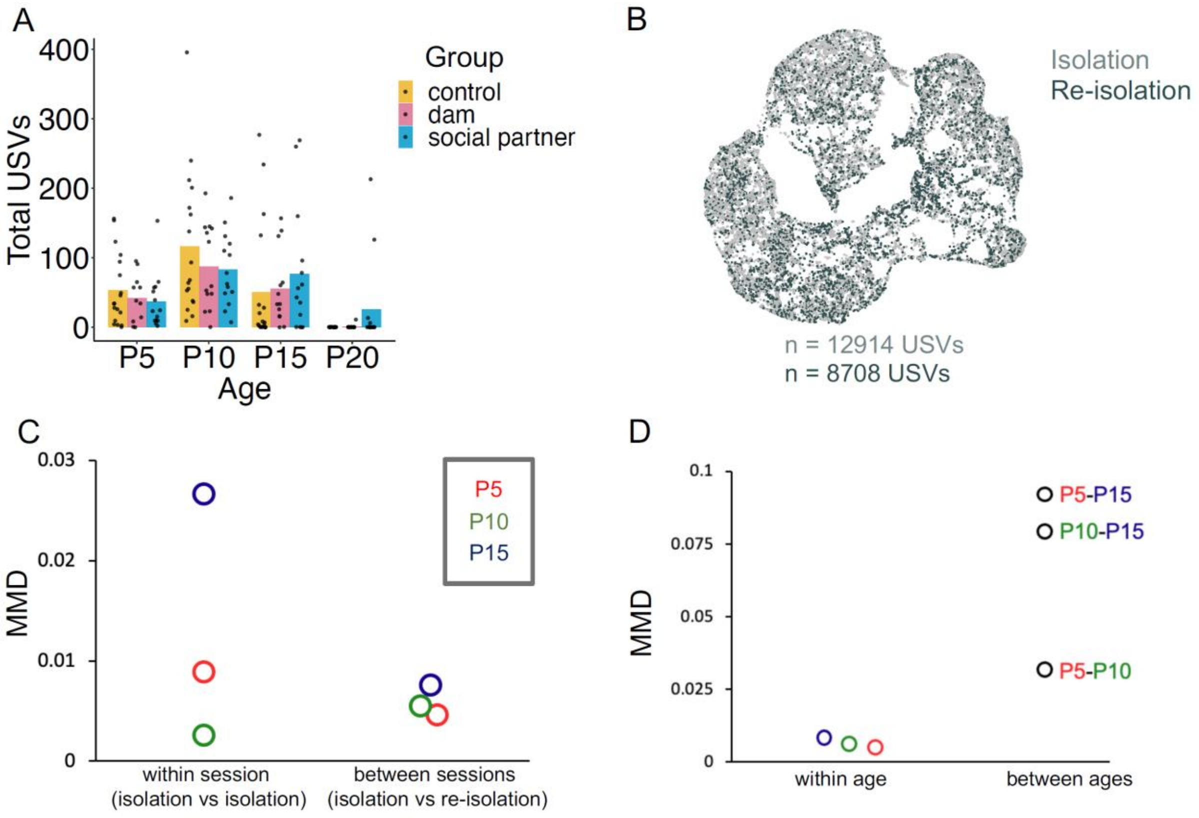

A
Total USVs
400
300
200
100
0
P5
P10
P15
P20
Age
Group
control
dam
social partner
B
Isolation
Re-isolation
n = 12914 USVs
n = 8708 USVs
C
MMD
0.03
0.02
0.01
0
P5
P10
P15
within session
(isolation vs isolation)
between sessions
(isolation vs re-isolation)
D
MMD
0.1
0.075
0.05
0.025
0
P5-P15
P10-P15
P5-P10
within age
between ages

**Figure S1. Further quantification of rates and acoustic features of USVs produced during isolation and re-isolation sessions.** (A) Number of USVs recorded during re-isolation sessions at P5, P10, P15 and P20. In the immediately preceding 5-minute social session, pups were recorded either with their dam, with a novel adult female, or with no social partner (control). No significant differences in rates of re-isolation USVs were found between social groups at any age ($p > 0.05$ for all within age comparisons). (B) UMAP projections of latent syllable representations of USVs produced by P5, P10, and P15 mice during isolation (gray points) and re-isolation sessions (black points). Points represent individual syllables and are closer to each other if acoustically more similar. (C) Maximum Mean Discrepancy (MMD) was calculated between distributions of latent syllable representations to generate two comparisons for each age: isolation vs. isolation (within session) and isolation vs. re-isolation (between sessions). To generate within-session comparisons, isolation sessions of different animals randomly assigned to one of two groups were compared. Comparisons with higher MMD values are more dissimilar. (D) MMD values were calculated between distributions of latent syllable representations to compare differences in acoustic features of isolation USVs within ages and between ages. Comparisons with higher MMD values are more dissimilar. Within age comparisons were generated by considering USVs produced during isolation sessions vs. USVs produced during re-isolation sessions and calculating MMD between these two distributions

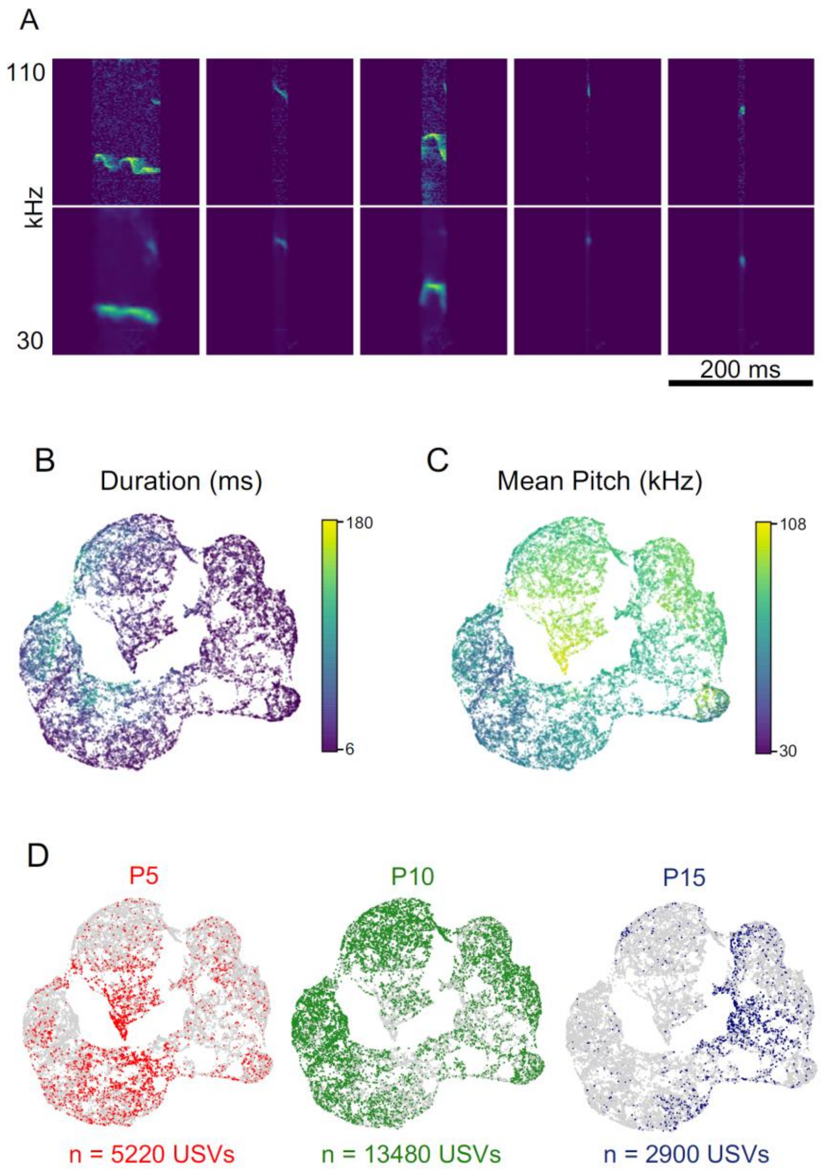

**Figure S2. UMAP projections of latent syllable representations and reconstructed spectrograms of isolation USVs produced by P5, P10, and P15 mice, color-coded by hand-picked acoustic features.** (A) Reconstructed spectrograms of representative syllables. (B) UMAP projections of syllable representations are color-coded by duration (ms). (C) UMAP projections of syllable representations are color-coded by mean pitch (kHz). (D) UMAP projections of syllable representations color-coded by age (P5, red; P10, green; P15, blue) are reproduced from Figure 1C for comparison.

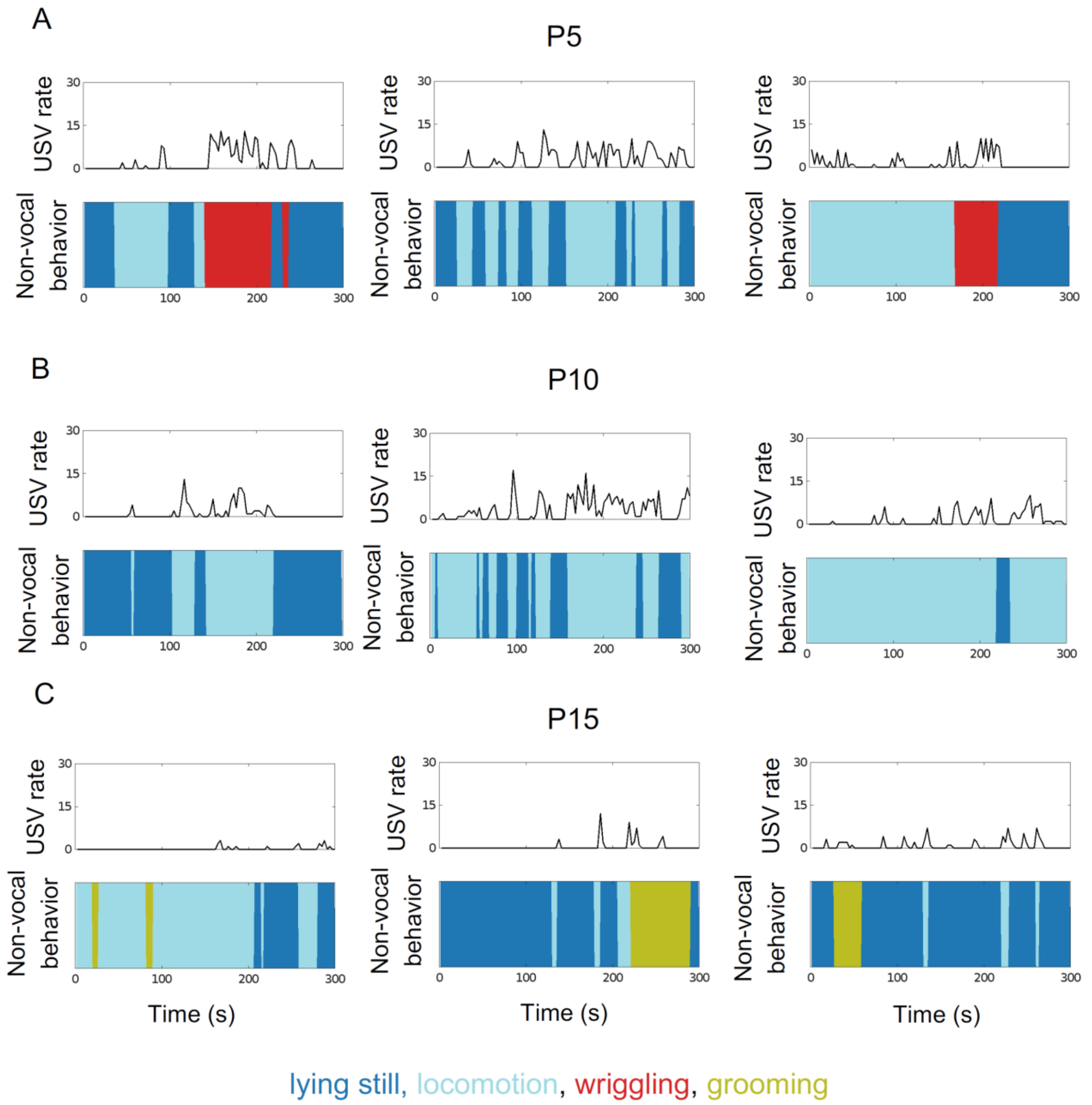

**Figure S3. Ethograms of USV production and non-vocal behavior.** Representative ethograms are shown for trials from P5 mice (A), P10 mice (B) and P15 mice (C). The top half of each plot shows USV rate over time (total USVs in each 3s-long bin), and the bottom half of each plot shows the occurrence of non-vocal behaviors across time (lying still, dark blue; locomotion, light blue; wriggling, red; grooming: chartreuse).

**Figure S4**

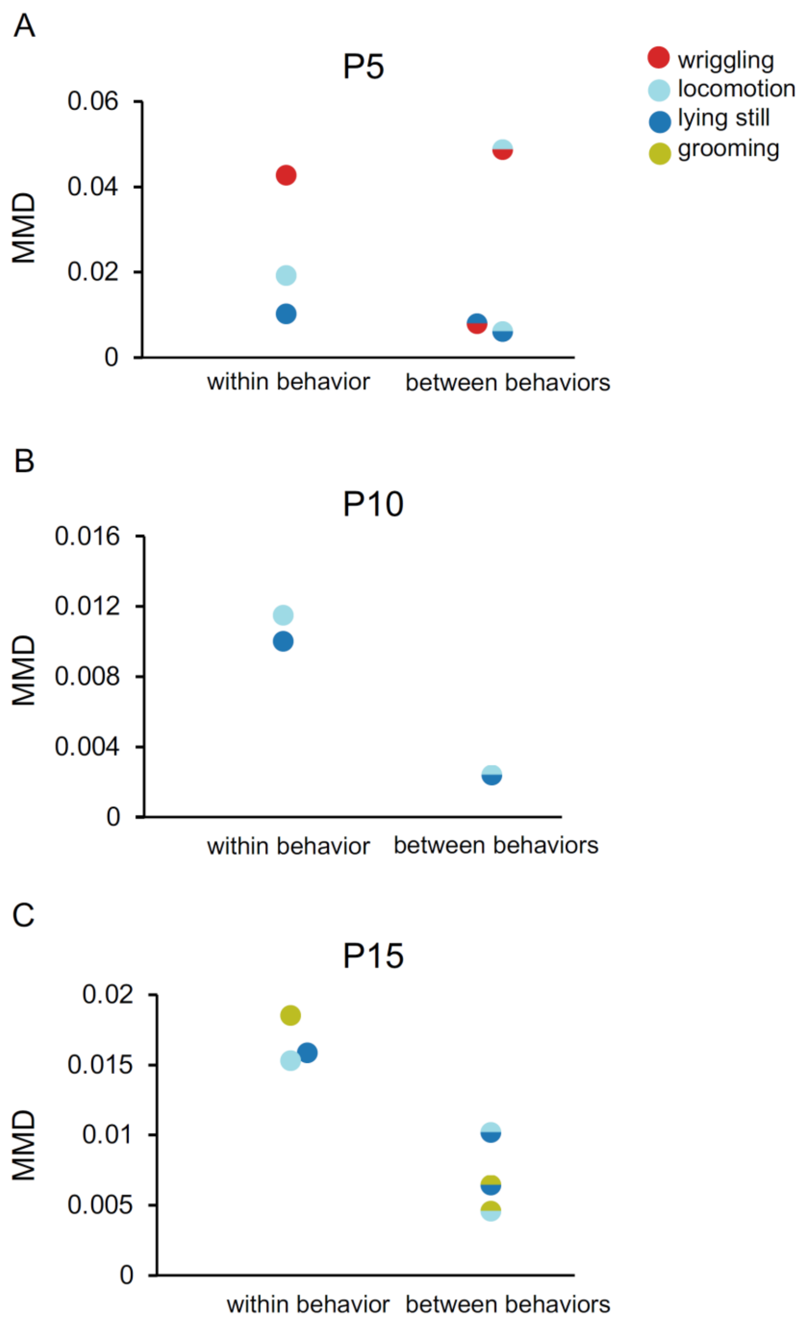
A
P5
0.06
0.04
0.02
0
MMD
within behavior
between behaviors
wriggling
locomotion
lying still
grooming
B
P10
0.016
0.012
0.008
0.004
0
MMD
within behavior
between behaviors
C
P15
0.02
0.015
0.01
0.005
0
MMD
within behavior
between behaviors

**Figure S4. Maximum Mean Discrepancy between distributions of latent representations of isolation USVs produced during different categories of non-vocal behavior.** (A) MMD values were calculated between distributions of latent syllable representations to compare differences in acoustic features of isolation USVs produced during different non-vocal behaviors in P5 pups. (B) Same as (A), for P10 USVs. (C) Same as (A), for P15 USVs. Comparisons with higher MMD values are more dissimilar.

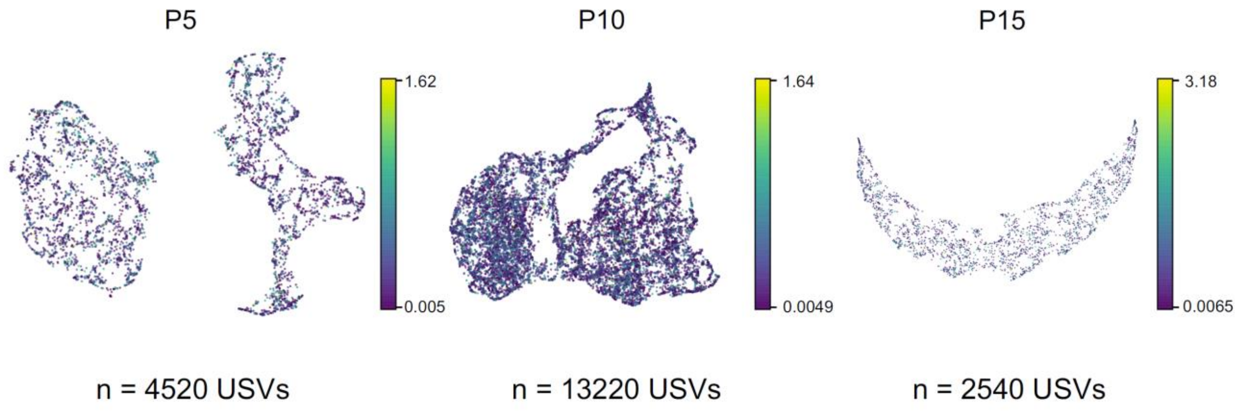

**Figure S5. Relationship of USV acoustic features to intensity of non-vocal movements.** UMAP projections of latent descriptions of USV syllables produced by P5 (left), P10 (middle) and P15 mice (right). Individual syllable representations are outlined in color according to the intensity of non-vocal movement that occurred while that syllable was produced. Please note that the by-age bokeh plots differ in shape from those that appear in Fig. 4B because the VAE was trained independently for these two analyses and generates different models and therefore different latent representations of USV syllables, even when trained on the same inputs.

**Movie S1. Example of USVs produced during wriggling.** Representative video (top) of a P5 pup producing USVs during wriggling. Bottom panel shows a spectrogram of USVs synchronized to the video, and the audio is pitch shifted (80 kHz to 5 kHz) to a range audible by humans.

**Movie S2. Example of USVs produced during locomotion.** Representative video (top) of a P10 pup producing USVs while engaged in locomotion. Bottom panel shows a spectrogram of USVs synchronized to the video, and the audio is pitch shifted (80 kHz to 5 kHz) to a range audible by humans.

**Movie S3. Example of USVs produced while lying still.** Representative video (top) of a P10 pup producing USVs while lying still. Bottom panel shows a spectrogram of USVs synchronized to the video, and the audio is pitch shifted (80 kHz to 5 kHz) to a range audible by humans.

**Movie S4. Example of USVs produced while grooming.** Representative video (top) of a P15 pup producing USVs during grooming. Bottom panel shows a spectrogram of USVs synchronized to the video, and the audio is pitch shifted (80 kHz to 5 kHz) to a range audible by humans.

**Table S1. Statistical summary.** Details of statistical analyses used in this study are presented.

| Figure | Comparison | Tests | Outcome | Notes |
|---|---|---|---|---|
| Fig. 1B | USV counts x age | One-way ANOVA; post-hoc t-tests with Bonferroni corrections | $p < 0.001$ (main effect of age) ** <br> $p < 0.001$ (P5 vs. P10) ** <br> $p = 0.31$ (P5 vs. P15) <br> $p < 0.001$ (P5 vs. P20) ** <br> $p < 0.001$ (P10 vs. P15) ** <br> $p < 0.001$ (P10 vs. P20) ** <br> $p = 0.16$ (P15 vs. P20) | |
| Fig. 1C, left | Syllable duration x age | Kruskal-Wallis, post-hoc Mann Whitney U tests with Bonferroni corrections | $p < 0.001$ (main effect of age) ** <br> $p < 0.001$ (P5 vs. P10) ** <br> $p < 0.001$ (P5 vs. P15) ** <br> $p < 0.001$ (P10 vs. P15) ** | P20 excluded; too few USVs |
| Fig. 1C, middle | Inter-syllable interval x age | Kruskal-Wallis, post-hoc Mann Whitney U tests with Bonferroni corrections | $p < 0.001$ (main effect of age) ** <br> $p < 0.001$ (P5 vs. P10) ** <br> $p < 0.001$ (P5 vs. P15) ** <br> $p = 0.001$ (P10 vs. P15) ** | P20 excluded; too few USVs |
| Fig. 1C, right | Syllable mean pitch x age | Kruskal-Wallis, post-hoc Mann Whitney U tests with Bonferroni corrections | $p < 0.001$ (main effect of age) ** <br> $p < 0.001$ (P5 vs. P10) ** <br> $p = 0.12$ (P5 vs. P15) <br> $p = 1.00$ (P10 vs. P15) | P20 excluded; too few USVs |
| Fig. S1A | Re-isolation USV counts x age x group (control, social partner, or dam) | 2-way ANOVA | $p < 0.001$ (main effect of age) ** <br> $p = 0.92$ (main effect of group) <br> $p = 0.94$ (interaction effect between age vs. group) | |
| Fig. 2 | Proportion time spent performing non-vocal behaviors x age | 2-way ANOVA; post-hoc t-tests with Bonferroni corrections | $p = 1.00$ (main effect of age) <br> $p < 0.001$ (main effect of non-vocal behavior) ** <br> $p < 0.001$ (interaction effect between non-vocal behavior vs. age) ** <br> $p < 0.001$ (P5 vs. P10 wriggling) ** <br> $p < 0.001$ (P5 vs. P15 wriggling) ** <br> $p = 1.00$ (P10 vs. P15 wriggling) | |

| | | | | |
|---|---|---|---|---|
| | | | p < 0.001 (P5 vs. P10 locomotion) ** <br>p < 0.001 (P5 vs. P15 locomotion) ** <br>p = 0.07 (P10 vs. P15 locomotion)<br>p < 0.001 (P5 vs. P10 lying still) **<br>p < 0.001 (P5 vs. P15 lying still) **<br>p = 0.12 (P10 vs. P15 lying still)<br>p = 1.00 (P5 vs. P10 grooming)<br>p < 0.001 (P5 vs. P15 grooming) **<br>p < 0.001 (P10 vs. P15 grooming) ** | |
| Fig. 3A (left) | USVs produced per second of each behavior at P5 | Friedman, post-hoc paired Wilcoxon-signed-rank tests with Bonferroni corrections | p = 0.04 (main effect of non-vocal behavior) *<br>p = 0.01 (locomotion vs. lying still) *<br>p = 0.55 (locomotion vs. wriggling)<br>p = 0.01 (lying still vs. wriggling) * | No grooming USVs at P5 |
| Fig. 3A (middle) | USVs produced per second of each behavior at P10 | Friedman, post-hoc paired Wilcoxon-signed-rank tests with Bonferroni corrections | p < 0.001 (main effect of non-vocal behavior) **<br>p < 0.001 (locomotion vs. lying still) **<br>p < 0.001 (locomotion vs. wriggling) **<br>p < 0.001 (lying still vs. wriggling) ** | No grooming USVs at P10 |
| Fig. 3A (right) | USVs produced per second of each behavior at P15 | Friedman, post-hoc paired Wilcoxon-signed-rank tests with Bonferroni corrections | p < 0.001 (main effect of non-vocal behavior) **<br>p < 0.001 (locomotion vs. lying still) **<br>p < 0.001 (locomotion vs. grooming) **<br>p = 0.04 (lying still vs. grooming) * | No wriggling USVs at P15 |
| Fig. 4B (left) | USV syllable duration x non-vocal behavior at P5 | Kruskal-Wallis, post-hoc Mann Whitney U tests with Bonferroni corrections | p < 0.001 (main effect of non-vocal behavior) **<br>p = 0.001 (locomotion vs. lying still) **<br>p = 0.001 (wriggling vs. lying still) **<br>p < 0.001 (wriggling vs. locomotion) ** | No grooming USVs at P5 |

| | | | | |
|---|---|---|---|---|
| Fig. 4B (middle) | USV syllable duration x non-vocal behavior at P10 | Kruskal-Wallis, post-hoc Mann Whitney U tests with Bonferroni corrections | $p = 0.004$ (main effect of non-vocal behavior) **<br>$p = 0.003$ (locomotion vs. lying still) ** | Very few wriggling USVs at P10 (n=41), excluded from analysis; no grooming USVs at P10 |
| Fig. 4B (right) | USV syllable duration x non-vocal behavior at P15 | Kruskal-Wallis, post-hoc Mann Whitney U tests with Bonferroni corrections | $p < 0.001$ (main effect of non-vocal behavior) **<br>$p < 0.001$ (locomotion vs. lying still) **<br>$p = 0.17$ (grooming vs. lying still)<br>$p = 0.38$ (grooming vs. locomotion) | No wriggling USVs at P15 |
| Fig. 4C (left) | USV syllable mean pitch x non-vocal behavior at P5 | Kruskal-Wallis, post-hoc Mann Whitney U tests with Bonferroni corrections | $p = 0.03$ (main effect of non-vocal behavior) *<br>$p = 0.14$ (locomotion vs. lying still)<br>$p = 1.00$ (wriggling vs. lying still)<br>$p = 0.04$ (wriggling vs. locomotion) * | No grooming USVs at P5 |
| Fig. 4C (middle) | USV syllable mean pitch x non-vocal behavior at P10 | Kruskal-Wallis, post-hoc Mann Whitney U tests with Bonferroni corrections | $p < 0.001$ (main effect of non-vocal behavior) **<br>$p < 0.001$ (locomotion vs. lying still) ** | Very few wriggling USVs at P10 (41), excluded from analysis; no grooming USVs at P10 |
| Fig. 4C (right) | USV syllable mean pitch x non-vocal behavior at P15 | Kruskal-Wallis, post-hoc Mann Whitney U tests with Bonferroni corrections | $p < 0.001$ (main effect of non-vocal behavior) **<br>$p < 0.001$ (locomotion vs. lying still) **<br>$p = 0.03$ (grooming vs. lying still) *<br>$p < 0.001$ (grooming vs. locomotion) ** | No wriggling USVs at P15 |

| | | | |
|---|---|---|---|
| Fig. 5 | Matched and shuffled comparisons of mean pooled covariance coefficients across and within ages | 2-way ANOVA with repeated measures on one factor; post-hoc t-tests with Bonferroni corrections | $p < 0.001$ (main effect of age) **<br>$p < 0.001$ (main effect of comparison type) **<br>$p < 0.001$ (interaction effect between comparison type vs. age) **<br>$p = 0.01$ (P5 vs. P10 matched) *<br>$p = 0.003$ (P5 vs. P15 matched) **<br>$p < 0.001$ (P10 vs. P15 matched) **<br>$p = 1.00$ (P5 vs. P10 shuffled)<br>$p = 1.00$ (P5 vs. P15 shuffled)<br>$p = 0.78$ (P10 vs. P15 shuffled) |
